# Cross-disorder GWAS meta-analysis for Attention Deficit/Hyperactivity Disorder, Autism Spectrum Disorder, Obsessive Compulsive Disorder, and Tourette Syndrome

**DOI:** 10.1101/770222

**Authors:** Zhiyu Yang, Hanrui Wu, Phil H. Lee, Fotis Tsetsos, Lea K. Davis, Dongmei Yu, Sang Hong Lee, Søren Dalsgaard, Jan Haavik, Csaba Barta, Tetyana Zayats, Valsamma Eapen, Naomi R. Wray, Bernie Devlin, Mark Daly, Benjamin Neale, Anders D. Børglum, James J. Crowley, Jeremiah Scharf, Carol A. Mathews, Stephen V. Faraone, Barbara Franke, Manuel Mattheisen, Jordan W. Smoller, Peristera Paschou

**Affiliations:** Department of Biological Sciences, Purdue University, West Lafayette, Indiana; Psychiatric and Neurodevelopmental Genetics Unit, Center for Genomic Medicine, Department of Psychiatry, Harvard Medical School, Massachusetts General Hospital, Boston, Massachusetts; Department of Molecular Biology and Genetics, Democritus University of Thrace, Alexandroupoli, Greece; the Division of Genetic Medicine, Vanderbilt Genetics Institute, Vanderbilt University Medical Center, Nashville, Tennessean; Queensland Brain Institute, University of Queensland, Brisbane, Queensland, 4072, Australia; Australian Centre for Precision Health, University of South Australia Cancer Research Institute, University of South Australia, Adelaide, South Australia, 5000, Australia; Lundbeck Foundation Initiative for Integrative Psychiatric Research, iPSYCH, Aarhus, Denmark; National Centre for Register-based Research, Aarhus University, Aarhus, Denmark; Hospital of Telemark, Kragerø, Norway; K.G. Jebsen Centre for Neuropsychiatric Disorders, Department of Biomedicine, University of Bergen, Norway; Division of Psychiatry, Haukeland University Hospital, Bergen, Norway; Institute of Medical Chemistry, Molecular Biology and Pathobiochemistry, Semmelweis University, Budapest, Hungary; Analytic Translational Genetics Unit, Massachusetts General Hospital and Harvard Medical School, Boston, Massachusetts; Academic Unit of Child Psychiatry South West Sydney, School of Psychiatry, University of New South Wales, Sydney, NSW, Australia; Institute for Molecular Bioscience, University of Queensland, Brisbane, Australia; Department of Psychiatry, University of Pittsburgh School of Medicine, Pittsburgh, Pennsylvania; Stanley Center for Psychiatric Research, Broad Institute, Cambridge, Massachusetts; Analytic and Translational Genetics Unit, Massachusetts General Hospital, Boston, Massachusetts; Medical and Population Genetics, Broad Institute, Cambridge, Massachusetts; Department of Biomedicine - Human Genetics, Aarhus University, Aarhus, Denmark; Center for Integrative Sequencing (iSEQ), Aarhus University, Aarhus, Denmark; Center for Genomics and Personalized Medicine, Aarhus, Denmark; Departments of Genetics and Psychiatry, University of North Carolina at Chapel Hill; Department of Psychiatry & Genetics Institute, University of Florida, Gainesville; Departments of Psychiatry and of Neuroscience and Physiology, SUNY Upstate Medical University, Syracuse, New York; Department of Human Genetics, Radboud University Medical Center, Nijmegen, The Netherlands; Department of Psychiatry, Radboud University Medical Center, Nijmegen, The Netherlands; Donders Institute for Brain, Cognition and Behaviour, Radboud University, Nijmegen, The Netherlands; Department of Psychiatry, Psychosomatics and Psychotherapy, Center of Mental Health, University Hospital Wuerzburg, Wuerzburg, Germany; Department of Biomedicine, Aarhus University, Aarhus, Denmark; Department of Clinical Neuroscience, Centre for Psychiatry Research, Karolinska Institutet, Stockholm, Sweden

**Keywords:** ADHD, ASD, OCD, Tourette Syndrome, cross-disorder genetic analysis, GWAS meta-analysis

## Abstract

Attention Deficit/Hyperactivity Disorder (ADHD), Autism Spectrum Disorder (ASD), Obsessive-Compulsive Disorder (OCD), and Tourette Syndrome (TS) are among the most prevalent neurodevelopmental psychiatric disorders of childhood and adolescence. High comorbidity rates across these four disorders point toward a common etiological thread that could be connecting them across the repetitive behaviors-impulsivity-compulsivity continuum. Aiming to uncover the shared genetic basis across ADHD, ASD, OCD, and TS, we undertake a systematic cross-disorder meta-analysis, integrating summary statistics from all currently available genome-wide association studies (GWAS) for these disorders, as made available by the Psychiatric Genomics Consortium (PGC) and the Lundbeck Foundation Initiative for Integrative Psychiatric Research (iPSYCH). We present analysis of a combined dataset of 93,294 individuals, across 6,788,510 markers and investigate associations on the single-nucleotide polymorphism (SNP), gene and pathway levels across all four disorders but also pairwise. In the ADHD-ASD-OCD-TS cross disorder GWAS meta-analysis, we uncover in total 297 genomewide significant variants from six LD (linkage disequilibrium) -independent genomic risk regions. Out of these genomewide significant association results, 199 SNPs, that map onto four genomic regions, show high posterior probability for association with at least three of the studied disorders (m-value>0.9). Gene-based GWAS meta-analysis across ADHD, ASD, OCD, and TS identified 21 genes significantly associated under Bonferroni correction. Out of those, 15 could not be identified as significantly associated based on the individual disorder GWAS dataset, indicating increased power in the cross-disorder comparisons. Cross-disorder tissue-specificity analysis implicates the Hypothalamus-Pituitary-Adrenal axis (stress response) as possibly underlying shared pathophysiology across ADHD, ASD, OCD, and TS. Our work highlights genetic variants and genes that may contribute to overlapping neurobiology across the four studied disorders and highlights the value of re-defining the framework for the study across this spectrum of highly comorbid disorders, by using transdiagnostic approaches.

## Introduction

Attention Deficit/Hyperactivity Disorder (ADHD, 5.3%), Autism Spectrum Disorder (ASD, 1.4%), Obsessive Compulsive Disorder (OCD, 1-3%), and Tourette Syndrome (TS, 0.6-1%) are among the most prevalent developmental psychiatric disorders of childhood and adolescence (1–4). Although each of these disorders defines a distinct DSM diagnostic category, they are highly comorbid. For instance, up to 55 % of TS patients also present with ADHD symptoms, 50% have OCD, and up to 20% present with ASD (5–7). The high comorbidity rates across these disorders lend support to the hypothesis of a common etiological thread that connects them across an impulsivity-compulsivity spectrum (8). Transdiagnostic approaches may shed light on the biology underlying this symptom continuum and hold the promise to identify targets for the development of personalized treatments that are still lacking.

ADHD, ASD, OCD, and TS all have a complex and highly heterogeneous genetic architecture with both common and rare genetic variants contributing to their etiology (4,9–12). Consequently, identifying and confirming genetic susceptibility factors has been challenging, demanding large samples for initial discovery and even larger samples for replication. Over the past few years, twelve genome-wide significant loci have been identified for ADHD (13), and five genome-wide significant loci were described for ASD (14,15). For OCD no genome-wide significant loci have been detected to date (16), while one genome-wide significant locus was recently reported for TS (17).

Based on the hypothesis for a shared etiology across ADHD, ASD, OCD, and TS (18,19), so far, several cross-disorder analyses have evaluated the genetic overlap across these disorders revealing broad genetic correlations (20–25). Most recently, the Psychiatric Genomics Consortium (PGC) presented a meta-analysis of GWAS across eight common psychiatric disorders including ADHD, ASD, OCD, and TS, analyzed jointly with GWAS data for anorexia nervosa (AN), bipolar disorder (BD), major depression (MD), and schizophrenia (SZ) (20). The study reports significant genetic correlations for most pairs of studied disorders, suggesting a complex, higher-order genetic structure underlying psychopathology. Exploratory factor analysis revealed three correlated factors, which together explained 51% of the genetic variation in the eight studied neuropsychiatric disorders. Early-onset disorders including ADHD, ASD, and, TS fell in one of the three identified factors (together with MD) while TS was also found in another factor together with compulsive disorders including OCD and AN. Variant-level analyses with all eight disorders analyzed jointly, supported the existence of substantial pleiotropy, with nearly 75% of the 146 genome-wide significant Single Nucleotide Polymorphisms (SNPs) influencing more than one of the eight examined disorders. Two loci among the ones that were found significant from the eight-disorder analysis were also reported with high confidence association for the four disorders that are the focus of the current study (implicating genes *DCC* and *RBFOX1*). These results further support the existence of shared neurobiology across traditional diagnostic boundaries revealing genetic loci that may have a pleiotropic effect across multiple psychiatric disorders. Thus studies that focus on cross-disorder etiology in finer detail are warranted.

Motivated by the high comorbidity rates and existing hypotheses for shared etiology across ADHD, ASD, OCD, and TS we focus on cross-disorder GWAS meta-analysis among these specific phenotypes. We integrate all currently available genome-wide data for these disorders and perform systematic cross-disorder meta-analyses seeking to identify shared and divergent genetic factors across four disorders that are often observed comorbid in childhood and adolescence. Our work highlights variants and genes that may contribute to neurobiology across the impulsivity-compulsivity spectrum of phenotypes.

## Methods

### Data sources

Analyses were conducted using summary statistics from GWAS for ADHD, ASD, OCD, and TS as made available by the PGC. For ADHD, samples were collected by iPSYCH and PGC, with most of the samples genotyped using the Illumina PsychArray. Only samples of European ancestry were included in our analyses, comprising 19,099 cases and 34,194 ancestry-matched controls. In total, 8,047,421 variants overlapping across all cohorts after imputation were analyzed (13).

For ASD, we acquired the summary statistics of 18,382 cases and 27,969 ancestry-matched controls of European ancestry collected by iPSYCH and PGC. Most of the samples were genotyped with the Illumina PsychChip. After meta-analysis, 9,112,387 variants overlapping across sample sources were available (15).

For OCD, we used results from a meta-analysis of GWAS from two consortia: International Obsessive Compulsive Disorder Foundation Genetics Collaborative (IOCDF-GC) (26) and OCD Collaborative Genetics Association Studies (OCGAS) (27), which led to a total of 2,688 affected samples and 7,037 ancestry-matched controls from Europe. Samples were genotyped with multiple different Illumina’s BeadChip arrays. After meta-analysis, 8,409,517 variants were found overlap and used for our study (16).

For TS, we combined results from the first GWAS on TS, conducted by Scharf et al. (28) and newly collected cases and controls. In total, 4,232 cases and 8,283 ancestry-matched controls were used for the analysis, which resulted in 8,868,895 variants overlapping in the meta-analysis. These summary statistics correspond to the GWAS carried out by Yu et al. (17), without samples from the Tic Genetic Consortium.

For all data obtained from the PGC, Ricopili pipeline (https://github.com/Ripkelab/ricopili/wiki) or comparable quality controls were carried out.

### Linkage disequilibrium (LD)-score regression to estimate genetic correlation across disorders

LD-score regression analysis was carried out using the LDSC package (29). Only common SNPs (MAF > 0.01) with an imputation quality (INFO) score > 0.9 and matched with the provided HapMap3 SNPs reference were analyzed. LD scores estimated for the European samples from the 1000 Genomes phase 3 (30) were used as both the independent variable and the weight for the regression.

### GWAS meta-analysis

To investigate the genetic variants underlying the observed overlap, cross-disorder meta-analysis was carried out for all four disorders combined, as well as for each individual pair of disorders that we found to be significantly correlated. SNP-based GWAS meta-analyses was performed using ASSET (31), which takes into account dependency across studies due to sample overlap (32). For each study, the variants’ effect sizes were measured by the logarithm of the odds ratio (OR). The possibility of inflation of results was investigated through observed λ as well as the sample size - corrected value λ_1000_. Variants with meta-analysis p-values below the genome-wide significance threshold (p < 5 x 10-8) were considered significant. To further highlight SNPs that contribute to risk across multiple phenotypes, we estimated the posterior probability of association (referred to as the m-value) with each disorder using a Bayesian statistical framework as implemented by MetaSoft (33). An m-value threshold of 0.9 has been recommended to predict with high confidence that a particular SNP is associated with a given disorder.

### Gene-based cross-disorder GWAS analysis

Gene-based cross-disorder GWAS analysis was carried out using the MAGMA plug-in on the FUMA GWAS annotation platform (34,35). For this analysis, variants were mapped onto genes based on their exact physical positions without extended windows and aggregated association p-values were calculated for each gene. Analysis was carried out under a SNP-wise (mean) model. Considering the sample composition, a European ancestry reference from 1000 Genomes phase 3 was used as the reference panel. Analysis was done with the summary statistics of each disorder individually as well as all meta-analysis results obtained. Significance thresholds were set applying Bonferroni correction for each analysis, corresponding to the number of genes being tested.

### Gene-property analysis for tissue specificity

To investigate phenotypic tissue specificity, a gene-property analysis testing for the relationship between tissue-specific gene expression and phenotype for associated genes was carried out using MAGMA for meta-analysis results with both GTEx v7 30 and 53 general tissue type expression atlas (36). Significant thresholds for these analyses were p-value < 1.67 x 10^-3^ and p-value < 9.43 x 10^-4^, respectively, under Bonferroni correction. The analysis was done for both the four-disorder and all the significantly correlated disorder pairs.

### Gene-set analysis

Gene-set analysis was also performed using MAGMA under a default competitive test. Gene sets and gene ontology (GO) terms tested were obtained from MsigDB v 6.1 (http://software.broadinstitute.org/gsea/msigdb), which contains 10,655 gene sets consistent across multiple sources. Bonferroni correction was applied to calculated association p-values to determine significance.

### Results annotation

SNP-based annotation and gene mapping were carried out for significant SNPs with ANNOVAR (37), including functional predictions for all significant non-synonymous mutations using SIFT (38) and PolyPhen-2 (39) plug-ins of ANNOVAR. Regional plots for the top-variants were created for 400 kb windows using the LocusZoom platform (40). For all significant results from our SNP-based and gene-based meta-analyses, we looked up previously reported associations in the GWAS catalog (41). Aggregate functional information and tissue expression levels of the genes were acquired from the GeneCards database (42), the GTEx Portal (43), and the Expression Atlas (44). Annotation of independent genomic risk loci from the FUMA GWAS platform was also adopted under parameters LD r^2^ < 0.6 for SNPs with association p < 5 x 10^-5^ and within 1000 kb away from the significant lead-SNP (p < 5 x 10^-8^). GO-annotation and the over-representation tests were performed using the R package ClusterProfiler v3.0.4 (45). Genes were mapped onto GO-terms based on org.Hs.eg.db (46). Enrichment of GO-terms was evaluated through a hypergeometric test (47). Network plotting was carried out using the built-in function of ClusterProfiler.

### Transcriptome-wide association study

Association between the studied disorders and gene expression levels in the brain was evaluated through summary-data-based Mendelian Randomization. The SMR software was used and analysis was performed for each individual disorder as well as using results from our GWAS meta-analyses (48). We used GWAS summary statistics for each studied disorder (as described above), the LD structure from from 1000 Genomes European reference panel and summary statistics from brain expression quantitative trait loci (eQTL) analysis (49), which quantified the effect of SNPs over gene expression levels in brain tissue (36,50). Only variants showing a consistent allele frequency (pairwise MAF difference between datasets no more than 0.20) across all three datasets (GWAS summary statistic, 1000 Genome reference, and eQTL summary statistic) were included in the analysis. All transcript probes with at least one cis-eQTL site showing p_eQTL_< 5 x 10^-8^ were taken into consideration. SNPs affecting the same probe with LD r^2^ > 0.9 or < 0.05 were pruned out from the analyses. Significance thresholds were based on Bonferroni correction for the number of probes tested.

To further verify that the effect of a probe on the trait was mediated by shared causal variants affecting both its expression and the trait rather than different variants in LD, we also carried out the HEterogeneity InDependent Instruments (HEIDI) test to evaluate the heterogeneity in the effect sizes of SNPs over trait and expression for each probe, evaluated as p_HEIDI_. As a default of the software, only SNPs with p_eQTL_ < 1.5654 x 10^-3^ were taken forward for this analysis. Up to top 20 independent SNPs in the cis-eQTL region were used for each tested probe to optimize the test power. A p_HEIDI_ > 0.05 indicates the existence of a shared cause underlying the expression level of a transcript probe and the trait, suggesting dysregulation of the transcript is functionally relevant to the trait.

## Results

### Genetic correlation across ADHD, ASD, OCD, and TS

First, we evaluated the extent of genetic overlap across ADHD, ASD, OCD, and TS using LD-score regression (Table 1). Results were in concordance with previous estimates (20). High genetic correlations were observed between all pairs of disorders, except for ASD and OCD. The highest genetic correlation was found between OCD and TS (rg = 0.3846, p = 0.0002), while a negative genetic correlation was observed between ADHD and OCD (rg = −0.1695, p = 0.0216). All regression intercepts for individual disorder heritability estimations were close to 1 (ADHD: 1.0334 (0.0101); ASD: 1.0084 (0.0096); OCD: 0.9928 (0.0068); TS: 1.0125 (0.0066)), indicating no signs of inflation.

**Table 1.** LD score regression analysis showing pairwise genetic correlation across ADHD, ASD, OCD, and TS. *#SNPs* = number of overlapping SNPs used in the analysis; *Rg* = genetic correlation; *SE, P* = standard error and *p-value* for *Rg; Intercept (SE)* = Intercept for genetic correlation and corresponding standard error.

|  | Disorder pairs |  |  |  |  |  |
| --- | --- | --- | --- | --- | --- | --- |
|  | ADHD/ASD | ADHD/OCD | ADHD/TS | ASD/OCD | ASD/TS | OCD/TS |
| #SNPs | 1042563 | 1030018 | 1062415 | 1012959 | 1044625 | 1100873 |
| $R_g$ | 0.3459 | -0.1695 | 0.2555 | 0.1185 | 0.177 | 0.3846 |
| SE | 0.0511 | 0.0738 | 0.06 | 0.0827 | 0.0638 | 0.1042 |
| P | 1.33E-11 | 0.0216 | 2.05E-05 | 0.1517 | 0.0055 | 0.0002 |
| Intercept | 0.3626 | 0.0082 | 0.0136 | -0.0003 | 0.0045 | 0.0728 |
| (SE) | (0.0073) | (0.006) | (0.0064) | (0.0063) | (0.0056) | (0.005) |

### Cross-disorder meta-analysis of ADHD, ASD, OCD, and TS GWAS

In order to identify genetic loci that are associated with patient phenotype across the spectrum of ADHD, ASD, OCD, and TS, we performed meta-analysis of available GWAS datasets. Combining all four datasets described above, 93,294 non-overlapping samples (including 51,311 controls) were available for analysis. After all filtering and cross-datasets alignment (as described in methods), 6,788,510 variants were retained. We found λ_ADHD-ASD-OCD-TS_ = 1.185, corrected λ_1000_ = 1.0040, showing little sign of inflation (Figure 1). We observed 297 genome-wide significant variants mapping onto six LD-independent genomic risk regions (Figure 1, Table S1, S2). 177 of the significant SNPs showed same direction of effect in all four disorders. Although we did not identify any SNP with m-value > 0.9 in all four disorders, all significant results had m-value > 0.9 in at least two of the disorders, and 199 were associated with m-value > 0.9 in three of the four disorders (Table S1). These 199 SNPs correspond to four genomic regions as shown in Table 2. The top result of the ADHD-ASD-OCD-TS GWAS meta-analysis was rs2144782 on chromosome 20, gene *KIZ/KIZ-AS1.* Three of the studied disorders (ADHD, ASD, and OCD) showed strong probability of association with this particular SNP (p = 4.18 x 10^-10^, m_ADHD_ = 1, m_ASD_ = 1, m_OCD_ = 0.911, m_TS_ = 0.672). Furthermore, two significantly associated SNPs in this region had m-values > in all four disorders indicating high probability for a cross-disorder effect (rs12625304, p = 8.12 x 10^-9^, m_ADHD_ = 0.986, m_ASD_ = 1, m_OCD_ = 0.918, m_TS_ = 0.881 and rs57080033, p = 2.90 x 10^-8^, m_ADHD_ = 0.999, m_ASD_ = 1, m_OCD_ = 0.83, m_TS_ = 0.911). This top region on chromosome 20 has also been previously highlighted by the ASD individual GWAS as well as results from the Cross-Disorder GWAS on eight different psychiatric disorders, although it was not reported among the most broadly pleiotropic ones (SNP rs6047287, p = 2.72 x 10^-10^) (Cross-Disorder Group of the Psychiatric Genomics Consortium et al., 2019).

**Figure 1.**
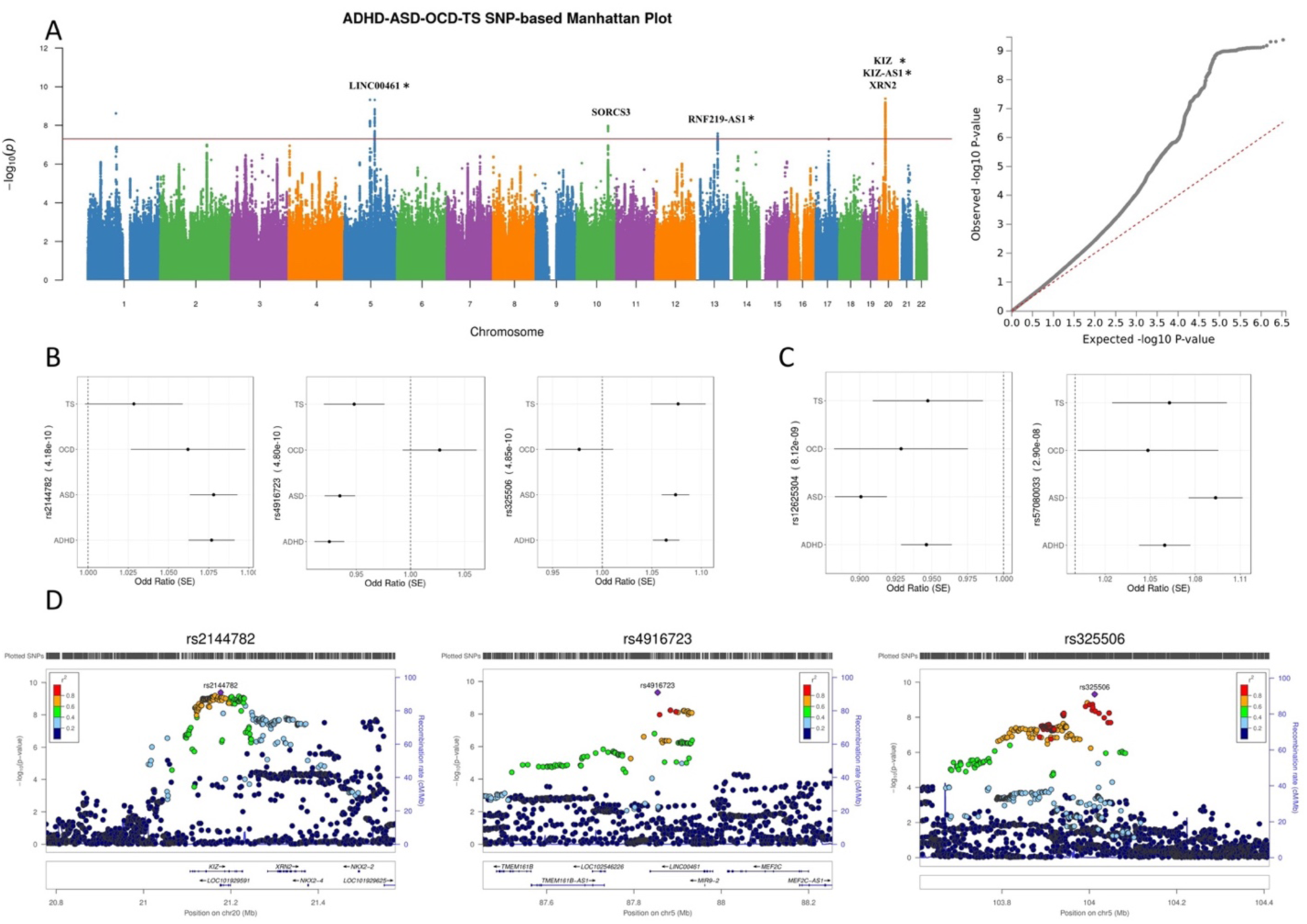
Cross-disorder ADHD-ASD-OCD-TS GWAS SNP-based meta-analysis. A. Manhattan plot and QQ plot for ADHD-ASD-OCD-TS GWAS meta-analysis. Genes harboring genome-wide significant SNPs are shown. An asterisk (*) indicates genes hosting SNPs with m-value > 0.9 in at least three disorders.; B. Forest plots for the top three SNPs; C. Forest plots for SNPs with m-value > 0.8 in all studied disorders; D. Regional plot for top three independent genetic loci.

**Table 2.** Cross-disorder ADHD-ASD-OCD-TS GWAS meta-analysis: Genetic risk loci hosting SNPs with m-value > 0.9 in at least three disorders (SNPs showing m-value>0.9 in at least 3 disorders that has the lowest P value in each region are shown).

| SNP | CHR | BP | P | Region | Gene(s) | ADHD m-value | ASD m-value | OCD m-value | TS m-value |
| --- | --- | --- | --- | --- | --- | --- | --- | --- | --- |
| rs2144782 | 20 | 21176604 | 4.18E-10 | 20p11.23 | KIZ-AS1, KIZ | 1 | 1 | 0.911 | 0.672 |
| rs325506 | 5 | 104012303 | 4.85E-10 | 5q21.2 | Intergenic | 1 | 1 | 0.13 | 0.981 |
| rs324895 | 5 | 87913831 | 6.22E-09 | 5q14.3 | LINC00461 | 1 | 0.999 | 0.419 | 0.935 |
| rs7318041 | 13 | 78857224 | 2.64E-08 | 13q22.3 | RNF219-AS1 | 1 | 1 | 0.264 | 0.908 |

In order to investigate cross-disorder relationships in finer detail, we performed pairwise SNP-based GWAS meta-analysis for those pairs of disorders that were significantly correlated (Table 1). Across ADHD and ASD, 8,093,883 overlapping variants were analyzed (λ_ADHD-ASD_ = 1.195, λ_1000_ = 1.0052). In this case, 374 SNPs surpassed the significance threshold (Figure 2, Table S1). These corresponded to seven independent genomic risk regions, one of which has not been previously reported as associated with either disorder, (gene *MANBA,* Table S2). The top-result of the ADHD-ASD GWAS meta-analysis was rs1222063 (p = 7.61 x 10^-11^, m_ADHD_ = 1, m_ASD_ = 1), an intergenic SNP on chromosome 1, residing between *LOC102723661* (distance = 114,004 bp) and *LINC01787* (distance = 117,185 bp). All 374 variants shared the same direction of effect and had m-values > 0.9 for both disorders indicating high probability for cross-disorder effect.

**Figure 2.**
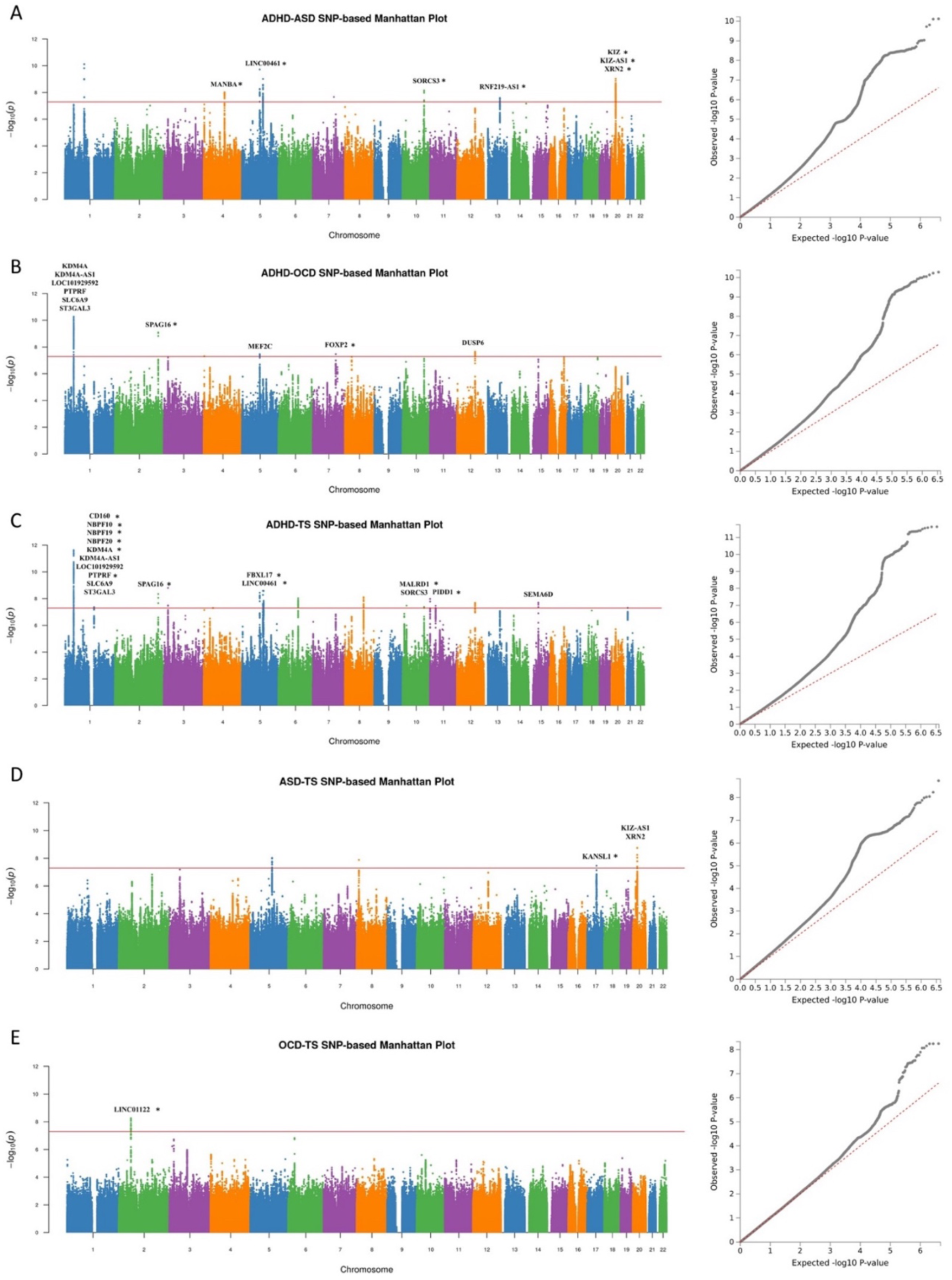
Manhattan plots and QQ plots for pairwise cross-disorder GWAS meta-analyses. An asterisk (*) indicates genes hosting SNPs with m-value > 0.9 in both disorders. A. ADHD-ASD GWAS meta-analysis; B. ADHD-OCD GWAS meta-analysis; C. ADHD-TS GWAS meta-analysis; D. ASD-TS GWAS meta-analysis; E. OCD-TS GWAS meta-analysis.

Across ADHD and OCD we analyzed 6,817,924 shared variants that passed the heterogeneity filter (λ_ADHD-OCD_ = 1.2020, λ _1000_ = 1.0071). 152 variants in six independent genomic regions were genome-wide significant, with one of them not identified in either ADHD or OCD individual study but showing m > 0.9 for both disorders here (Figure 2, Table S1, S2). As expected due to the negative genetic correlation uncovered between ADHD and OCD, 48 out of 152 significant SNPs showed opposite direction of effects in the two disorders and were driven by the ADHD GWAS (Table S1). Accordingly, only five of the variants showed m-value > 0.9 in both disorders, which can be mapped onto three independent risk loci (gene *SPAG16*, *FOXP2* and intergenic region on 4p16.3, table S1, S2). The top-association in the ADHD and OCD meta-analysis was found at rs113551349 (p = 5.25 x 10^-11^, m_ADHD_ = 1, m_OCD_ = 0.795) on chromosome 1, an intronic region of gene *SLC6A9*. On the other hand, the top SNP showing m-value > 0.9 in both disorders was rs9677504 (p = 8.04 x 10^-10^, m_ADHD_ = 1, m_OCD_ = 0.995) on chromosome 1, gene *SPAG16*.

Analysis across ADHD and TS was performed over 6,815,758 overlapping variants (λ_ADHD-TS_ = 1.2230, λ_1000_ = 1.0074), showing no sign of inflation. A total of 372 significant SNPs were identified (Figure 2, Table S1). The significant SNPs fell onto 16 independent genomic regions, six of which were novel (including variants at genes *CD160, NBPF10, NBPF19, NBPF20, FBXL17, MALRD1,* and *PIDD1*) (Table S2). All significant SNPs shared the same direction of effect, with almost half them (n=162) associated with m-values > 0.9 for both disorders studied and corresponding to as many as 12 different regions (Table S1, S2). The top-result was located on chromosome 1, rs112361411 and was possibly driven by the ADHD GWAS (p = 2.30 x 10^-12^, m_ADHD_ = 1, m_TS_ = 0.477). This variant resides within a non-coding RNA gene *LOC101929592* and was also detected in the individual GWAS on ADHD. On the other hand, the top result with high probability of shared effect across ADHD and TS was rs4660740 (p = 4.07 x 10^-12^, m_ADHD_ = 1, m_TS_ = 0.909) on chromosome 1, gene *KDM4A.* This locus has been previously associated with ADHD.

ASD and TS available GWAS datasets shared 7,499,503 SNPs. We found λ_ASD-TS_ = 1.1440, λ_1000_ = 1.0052, showing no sign of inflation. A total of 15 SNPs were genome-wide significant (Figure 2, Table S1). Four independent genomic risk regions were highlighted, one of which (at gene *KANSL1*) was not identified in either individual disorder GWAS (Table S2). The top-result, which had also been detected in the ASD GWAS, was rs1000177 (p = 1.79 x 10^-9^, m_ASD_ = 1, m_TS_ = 0.758) on chromosome 20. It is found in an intergenic region nearby *KIZ* (distance = 5,940 bp). As stated earlier here, this gene was also among the top-results of our four-disorder combined meta-analysis as well as the ADHD-ASD meta-analysis. All the significant SNPs had the same direction of effect in both disorders. Eight of the significant SNPs had m-values > 0.9 for both ASD and TS corresponding to two genomic risk regions (Table S1). Highest probability of shared effect for both ASD and TS was found for rs325506 (p = 9.42 x 10^-9^, m_ASD_ = 1, m_TS_ = 0.984) which lies on an intergenic region of chromosome 5.

Across the OCD and TS GWAS, 8,112,469 overlapping variants were available for analysis (λ_OCD-TS_ = 1.0020, λ_1000_ = 1.0002). No genome-wide significant SNP had been identified in the individual GWAS of OCD, and only a limited number of significant variants were revealed in the TS GWAS. In the meta-analysis across TS and OCD, we found 21 genome-wide significant variants (top-result rs140347666 (p = 5.64 x 10^-9^, m_OCD_ = 0.999, m_TS_ = 1); Figure 2, Table S1, S2); all significant SNPs were located in the same genomic risk region (chromosome 2, non-coding RNA gene *LINC01122*) and had the same direction of effect. All 21 SNP showed m-values > 0.9 for both TS and OCD, indicating high homogeneity across both disorders.

### Cross-disorder gene-based association and tissue-specificity analysis

We proceeded to perform gene-based and tissue-specificity analysis across ADHD, ASD, OCD, TS. With 18,450 protein-coding genes tested, the significance threshold for our gene-based analysis for the four-disorder combined meta-analysis (ADHD, ASD, OCD, TS) was 2.71 x 10^-6^ after Bonferroni correction. Our gene-based analysis highlighted 21 genes as significantly associated in the ADHD-ASD-OCD-TS meta-analysis. Out of those, 15 could not be identified as significant based on analysis of the individual disorder GWAS dataset, indicating increased power in the cross-disorder comparisons (Table S3). The top-result was *XRN2* (p = 2.08 x 10^-9^) on chromosome 20 (Figure 3, Table S3). The full list of significant results from our gene-based analysis for all four disorders as well as comparisons to results from the pairwise meta-analyses can be found in Figure 4 and Table S3. No gene was significant across all six meta-analyses while eight genes were found significant in three or more analyses (Table S3).

**Figure 3.**
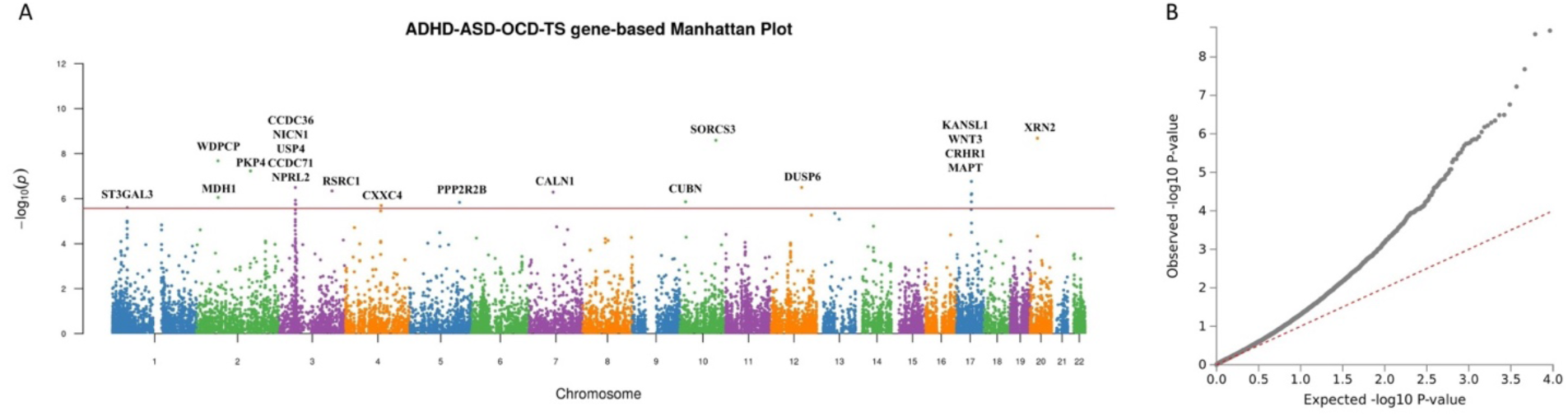
ADHD-ASD-OCD-TS cross-disorder gene-based GWAS meta-analysis. Manhattan plot (A) and QQ plot (B).

**Figure 4.**
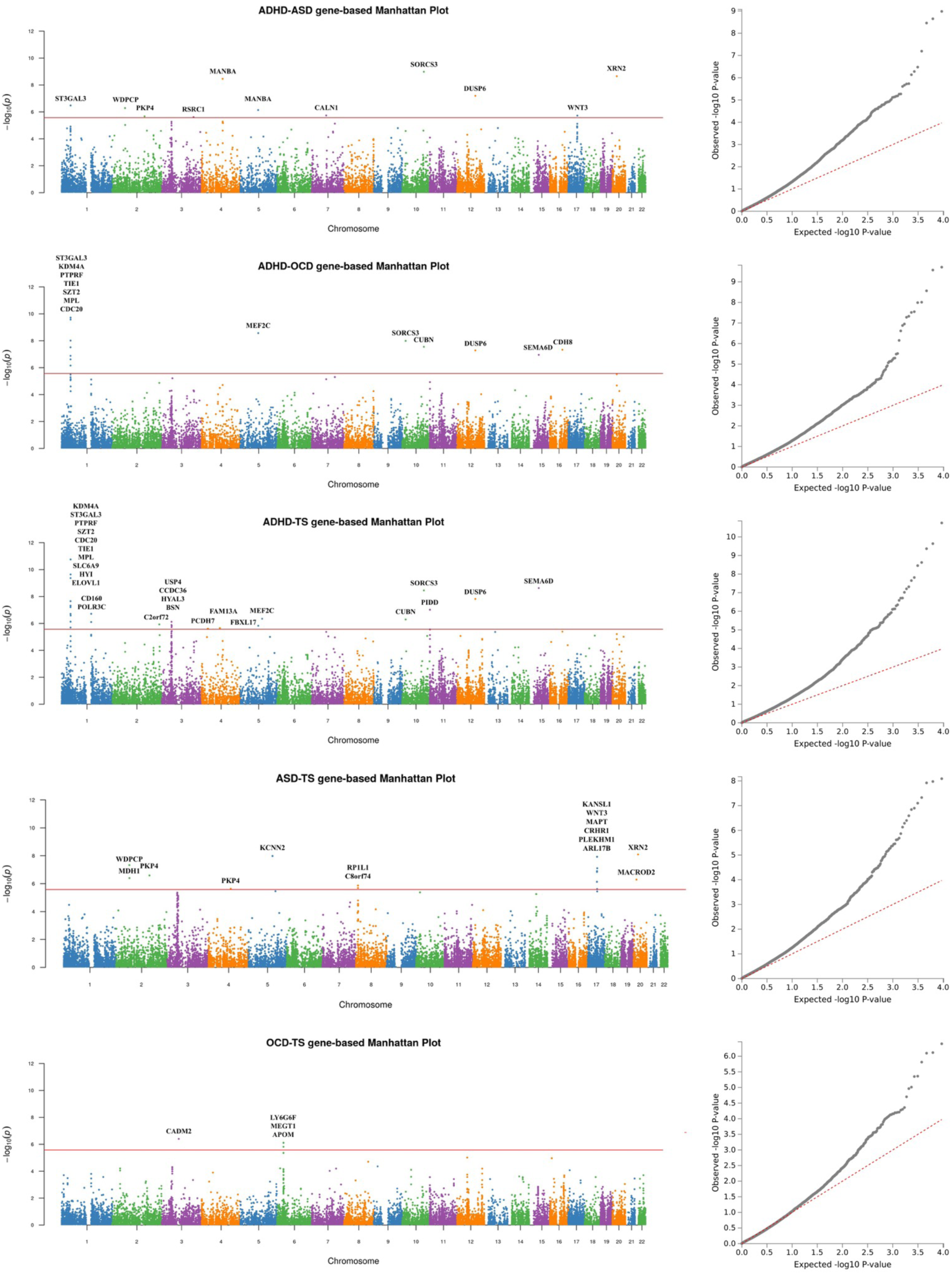
Manhattan plots for pairwise gene-based GWAS meta-analyses. A. ADHD-ASD gene-based analysis; B. ADHD-OCD gene-based analysis; C. ADHD-TS gene-based analysis; D. ASD-TS gene-based analysis; E. OCD-TS gene-based analysis.

In order to better visualize our results while investigating the pathways and interactions among the top risk genes across ADHD, OCD, ASD and TS, we constructed GO-based networks for the top 200 genes from each gene-based association analysis as well as genes annotated from the SNP-based GWAS meta-analyses. Results are shown in Figure 5 and Figure S1-S5. Pathways related to neuronal development, axonogenesis, and synaptic structure and organization were highlighted among the most significant in our analysis. These results were further strengthened by gene-property analyses, which showed enrichment of our top associated loci in genes expressed in brain tissues (Figure 6). In the tissue specificity analysis based on the ADHD-ASD-OCD-TS GWAS meta-analysis results, significant enrichment was found for genes expressed in the basal ganglia, hypothalamus, areas of the frontal cortex, and in the anterior cingulate cortex (Table S4). Additional brain regions implicated by our analysis were frontal cortex, cerebellum, amygdala, and hippocampus (Table S4). Furthermore, enrichment for genes expressed in the pituitary was also found among the top implicated regions from the ADHD-ASD-OCD-TS, ADHD-ASD and ADHD-TS GWAS meta-analyses. Intriguingly, besides showing enrichment of gene expression in the brain (including hypothalamus, Figure S6), and pituitary, the four-disorder combined cross-disorder analysis also identified an enrichment of gene expression in the adrenal gland thus pointing to involvement of the Hypothalamus-Pituitary-Adrenal gland axis across ADHD, ASD, OCD, and TS (Figure 6, S6 and Table S4).

**Figure 5.**
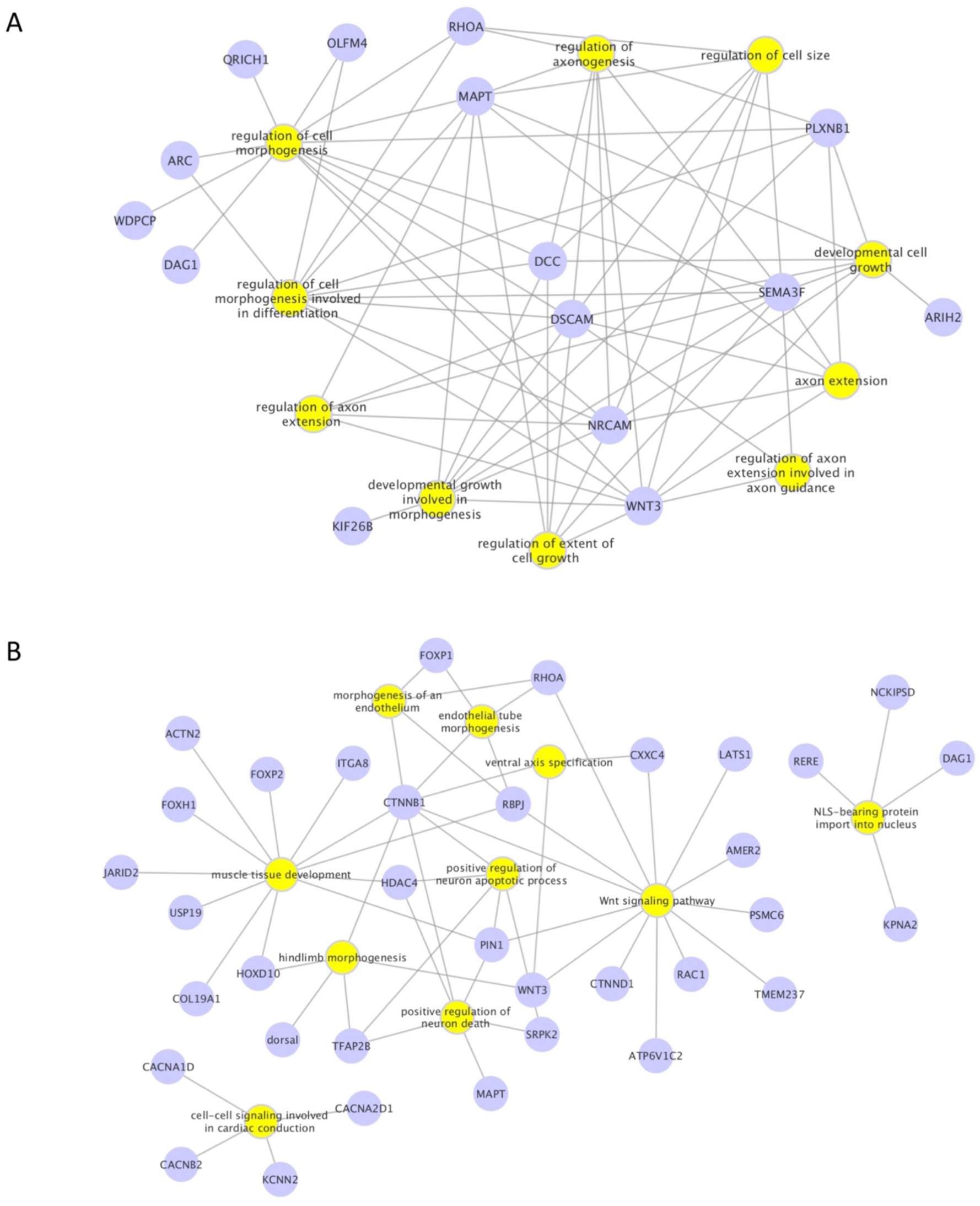
Gene networks for top genes in cross-disorder ADHD-ASD-OCD-TS GWAS meta-analysis. **A.** Top ten gene networks based on top 200 genes from SNP-based ADHD-ASD-OCD-TS meta-analyses results. **B.** Top ten gene networks based on top 200 genes from ADHD-ASD-OCD-TS gene-based analyses.

**Figure 6.**
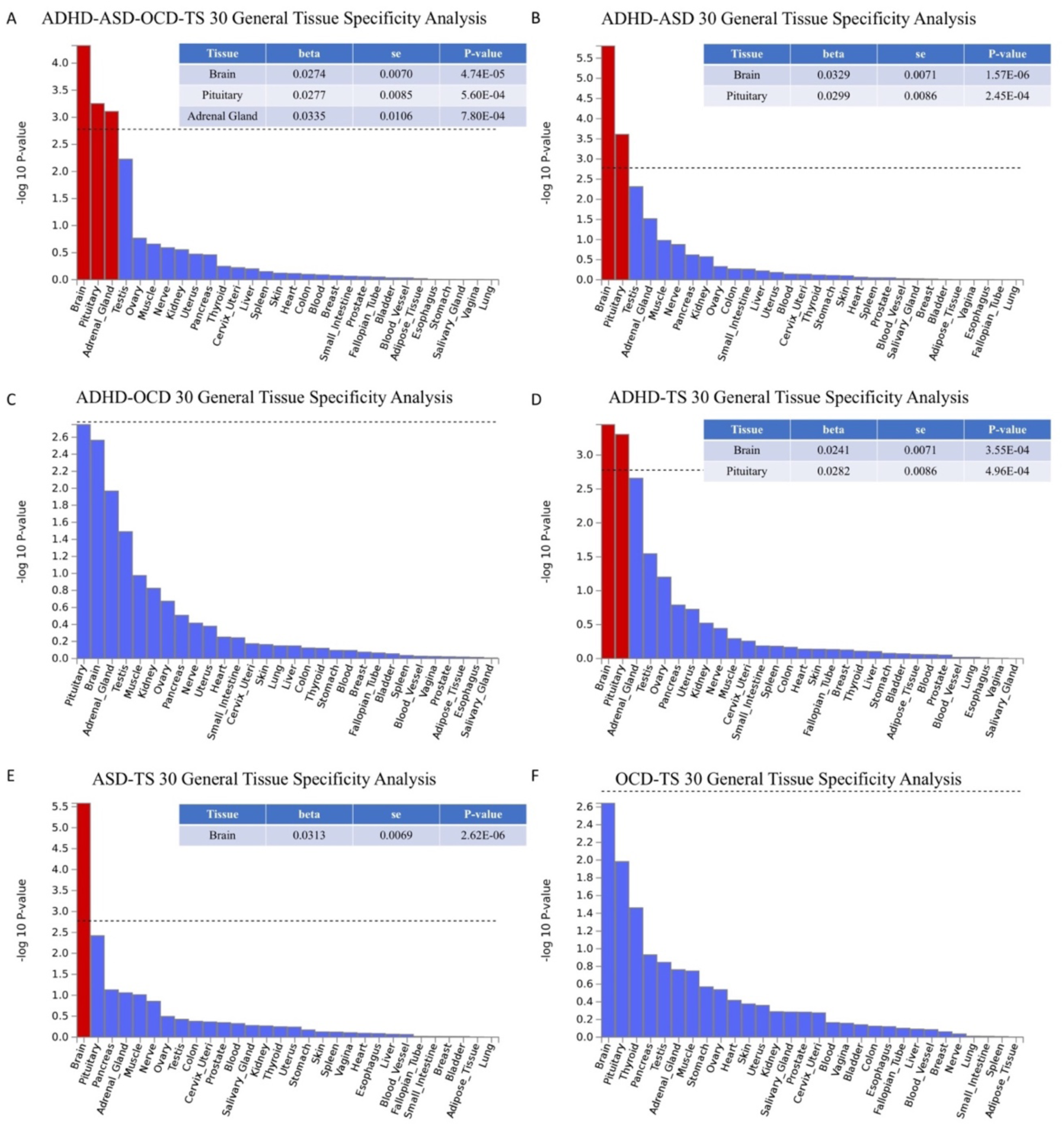
ADHD-ASD-OCD-TS cross-disorder tissue specificity analysis testing 30 general tissue types from GTEx v7 tissue expression atlas. Red bar indicates significant enrichment of gene expression in corresponding tissue under Bonferroni correction (p < 1.67 x 10-3). Panel on top right corner of each figure shows detailed statistics for significantly enriched tissue. A. ADHD-ASD-OCD-TS cross-disorder tissue specific expression enrichment; B. ADHD-ASD cross-disorder tissue specific expression enrichment; C. ADHD-OCD cross-disorder tissue specific expression enrichment; D. ADHD-TS cross-disorder tissue specific expression enrichment; E. ASD-TS cross-disorder tissue specific expression enrichment; F. OCD-TS cross-disorder tissue specific expression enrichment.

### Transcriptome-wide association results

Next, we incorporated eQTL information into our meta-analyses and performed transcriptome-wide association analyses, aiming to identify genes with expression levels associated across the studied disorders. For the ADHD-ASD-OCD-TS combined transcriptome-wide analysis, we used a total of 6,710,493 SNPs from the meta-analysis summary statistics which also presented in both 1000Genome and the eQTL summary statistics. In total 7,295 transcript probes were analyzed, corresponding to a significance threshold of p_SMR_ < 6.85 x 10^-6^. Under this threshold, five transcript probes were significant, all located on chromosome 17. Two of them also satisfied the pleiotropy hypothesis, which indicates the existence of variants with a shared effect over both gene expression level and trait. Among all significant transcripts, the top-result was from the *LRRC37A4P* probe (p_SMR_ = 2.20 x 10^-6^, p_HEIDI_ = 0.0843). This corresponds to the transcript of a pseudogene in region 17q21.3, localizing near *KANSL1*. The two significant, pleiotropic transcripts also turned out to be significant in the ASD-TS pairwise analysis we performed. Comparison of significant results obtained from all meta-analyses can be found in Table 3.

**Table 3.** Significant results from Transcriptome-wide association analyses. *pSMR = p-value* for transcriptome-wide association analysis; *PHEIDI = p-value* for heterogeneity in dependent instruments (HEIDI) test. *PHEIDI >* 0.05 indicates a pleiotropic SNP effect over both the trait and the probe expression.

| CHR | Gene | ADHD-ASD-OCD-TS |  |  |  | ADHD-TS |  |  |  | ASD-TS |  |  |  |
| --- | --- | --- | --- | --- | --- | --- | --- | --- | --- | --- | --- | --- | --- |
| | | Beta | SE | $p_{SMR}$ | $P_{HEIDI}$ | Beta | SE | $p_{SMR}$ | $P_{HEIDI}$ | Beta | SE | $p_{SMR}$ | $P_{HEIDI}$ |
| 11 | AP006621.1 |  |  |  |  | 0.0491 | 0.0109 | 6.67E-06 | 5.63E-01 |  |  |  |  |
| 11 | AP006621.5 |  |  |  |  | 0.0540 | 0.0110 | 9.73E-07 | 1.42E-01 |  |  |  |  |
| 17 | LRRC37A4P | -0.0333 | 0.0070 | 2.20E-06 | 8.43E-02 |  |  |  |  | -0.0561 | 0.0112 | 5.58E-07 | 1.68E-01 |
| 17 | RP11-707O23.5 | 0.0308 | 0.0067 | 3.90E-06 | 8.45E-02 |  |  |  |  | 0.0536 | 0.0106 | 3.83E-07 | 8.53E-02 |
| 17 | RP11-798G7.6 |  |  |  |  |  |  |  |  | 0.1239 | 0.0268 | 3.78E-06 | 4.87E-01 |
| 17 | KANSL1-AS1 |  |  |  |  |  |  |  |  | 0.0487 | 0.0104 | 3.09E-06 | 5.26E-02 |
| 17 | RP11-259G18.2 |  |  |  |  |  |  |  |  | 0.0512 | 0.0104 | 9.33E-07 | 1.48E-01 |
| 18 | MPPE1 |  |  |  |  |  |  |  |  | -0.0632 | 0.0139 | 5.89E-06 | 1.76E-01 |

We also ran transcriptome-wide association analyses for each individual disorder. After Bonferroni correction, significant results were only observed for ADHD and ASD. For ADHD, out of 7,295 probes tested, one was found significant (with p_SMR_ < 6.85 x 10^-6^ as significance threshold) without showing heterogeneity of effect over expression and trait (p_SMR_ = 6.25 x 10^-6^, p_HEIDI_ = 0.136). This probe tags *PIDD* on chromosome 11. For ASD (7,445 probes tested, with p_SMR_ < 6.72 x 10^-6^ as significance threshold), probes for *LRRC37A4P* (p_SMR_ = 2.22 x 10^-6^, p_HEIDI_ = 0.110) and *RP11-259G18.2* (p_SMR_ = 4.37 x 10^-6^, p_HEIDI_ = 0.277) on chromosome 17 were significant.

## Discussion

To our knowledge, this is the first study focusing on the detailed investigation of the shared genetic basis across ADHD, ASD, OCD and TS. Using available GWAS summary statistics for each individual disorder, we uncovered clues to the potential etiological overlap that may underlie the studied phenotypes. Pairwise LD-score regression results were concordant with previous PGC analysis (20), indicating high genetic correlations across the spectrum of studied phenotypes with two exceptions that have also been previously reported (20): the genetic correlation between ASD and OCD was not significant and there was a negative genetic correlation between ADHD and OCD. The lack of significant correlation between ADHD and OCD could potentially be due to the limited sample size of the OCD dataset. On the other hand, the negative genetic correlation between ADHD and OCD indicates that genetic variants operate in opposite directions in the development of these two disorders. From a clinical perspective, this is quite intuitive since ADHD and OCD may be thought of as lying at opposite extremes of the impulsivity-compulsivity continuum.

We successfully uncovered 297 genome-wide significant variants from six LD-independent genomic risk regions in the ADHD-ASD-OCD-TS GWAS cross-disorder meta-analysis. Interestingly, out of all the significant SNPs, as many as 261 could not be identified as significant at a genome-wide level based on any of the individual disorder GWAS datasets, indicating a boosted statistical power through the meta-analysis. All the significant results showed evidence of shared effect in at least two of the disorders, and 199 SNPs (mapping onto four genomic regions), showed high probability of association in three of the four disorders (as defined by m-value>0.9). The top-significant genomic risk locus showing also high probability for association across the studied disorders was region 20p11.23. Our top genomic risk locus had also been previously identified as significant in the individual ASD GWAS (15) and was also found significant in all our meta-analyses described here with the ASD GWAS involved. Furthermore, this region was also found significant in the eight psychiatric disorder meta-analysis carried out by the Cross-Disorder Group of the Psychiatric Genomics Consortium (20) with leading SNP rs6047287, p = 2.72 x 10^-10^). Gene *KIZ* hosts many of the top significant SNPs uncovered by our cross-disorder meta-analysis. It is widely expressed in diverse human tissues and is involved in cell mitosis. It is a core gene in the Polo-Like Kinase 1 (*PLK1*) pathway and is critical for regulating human spindle-pole formation (51,52). Previous work has shown that neurogenesis and the earliest phases of neuronal differentiation are compromised by spindle misorientation (51). Furthermore, *KIZ* is associated with systemic lupus erythematosus (53), and peanut allergy (54). These associations indicate a possible role of *KIZ* in immune response, a mechanism which is also implicated in TS etiology (55). Our ADHD-ASD-OCD-TS cross-disorder GWAS meta-analysis also highlighted the role of genomic regions 5q21.2 and 5q14.3 with significant effects contributed by ADHD, ASD, and TS. Duplication of the 5q21.2 region has been previously reported as a clinically significant copy number variation (CNV) in schizophrenia (56)

The top result from our four-disorder gene-based analysis was *XRN2* (ADHD/OCD/ASD/TS-p = 2.08 x 10^-9^, ADHD/ASD-p = 2.24 x 10^-9^, ASD/TS-p = 8.11 x 10^-9^). This gene is widely expressed in various human tissues and encodes an essential nuclear 5’→3’ exoRNase with many functions in the processing and regulation of RNA molecules (57). It has been previously associated with ASD (15,58). Furthermore, a recent large-scale transcriptome association study in ASD identified significant differential expression and splicing of *XRN2* in ASD, further suggesting a functional role of *XRN2* in this neurodevelopmental disorder (58).

*SORCS3* (ADHD/OCD/ASD/TS-p = 2.56 x 10^-9^, ADHD/ASD-p = 1.04 x 10^-9^, ADHD/OCD-p = 2.82 x 10^-8^, ADHD/TS-p = 3.51 x 10^-9^) was the second top-gene in the four-disorder gene-based meta-analysis. This gene encodes a member of the vacuolar protein sorting 10 (VPS10) receptor family, which controls intracellular protein signaling in neurons and regulates neuronal viability through many pathways (59). It is highly expressed in brain tissues (36), and it has been previously implicated in neurological disease including ADHD and ASD etiology (13,15). Multiple studies indicate a relationship between SORCS3 and the accumulation of amyloid, which is linked to Alzheimer disease (60,61). It is also associated with major depression in individuals of European descent (62). Moreover, its interaction with postsynaptic proteins, such as *PICK1*, indicates that the product of *SORCS3* regulates glutamate receptor function (63,64). As one of the major neurotransmitters in the human brain, the glutamate pathway has long been hypothesized to underlie abnormalities in ADHD, ASD, OCD, and TS and is a possible therapeutic target for these disorders (65–68).

Several of our top findings, including the *SORCS3* region on chromosome 10, the *KIZ* region on chromosome 20, and an intergenic region on chromosome 5, were also reported as genomewide significant in the recent GWAS meta-analysis of the PGC seeking pleiotropic loci across a broad spectrum of eight psychiatric disorders, including the four that are the focus of our study here (20). In fact, *SORCS3* and region chr5: 103,791,044-104,055,261 were reported among those genomic regions with broad cross-disorder association across the eight psychiatric disorders analyzed (20), indicating that they may have a general effect over neuropsychiatric disorders rather than specific for the disorders of high comorbidity and high prevalence in childhood and adolescence that we analyzed here. On the other hand, the two regions (region 18q21.2, gene *DCC* and region 16p13.3, gene *RBFOX1*) that were revealed as the most broadly pleiotropic in the PGC cross-disorder analysis were not found genomewide significant here, possibly due to differences in sample composition. Note however, that *DCC* is indeed revealed as one of the nodes in the gene-network analysis of the top 200 genes annotated from the SNP-based analysis for ADHD-ASD-OCD-TS, ADHD-OCD, ADHD-TS and network obtained from top 200 genes of the gene-based analysis for ADHD-ASD that we performed (Figure 5, Figure S2).

Among the top genes that we found associated in the ADHD-ASD-OCD-TS GWAS meta-analysis, we observed enrichment for genes expressed in the brain. Our results provide further support for the involvement of the basal ganglia across all disorders analyzed here. Dysfunction of the basal ganglia has been observed in all four studied disorders (69–72). Interestingly, our analyses also implicate the involvement of the hypothalamus-pituitary-adrenal (HPA) axis, in accordance with previous studies implicating this system in multiple childhood-onset psychiatric traits including ADHD and TS (73–77). The HPA axis plays a critical role in human stress response through the regulation of cortisol secretion (78). Low-cortisol responsivity to stress was proposed as a biomarker for certain types of ADHD, indicating a possibly altered HPA axis activity in this disorder (79). Altered cortisol levels among TS individuals have also been reported, with a negative correlation between evening cortisol and patients’ tic severity and higher cortisol levels in response to stress. Both observations further support the potential involvement of the HPA axis (80).

Gene ontology analyses highlighted the involvement of pathways related to neuronal development, cell adhesion, axonogenesis, synaptic structure, and synaptic organization. Furthermore, many of our top associated genes had been previously associated with intelligence (81), cognitive abilities (82), depressive symptoms (83), neuroticism (84), and neurological disorders such as Parkinson disease (85). Based on the observation that enriched gene expression was found in brain tissues for most of the disorder combinations we also investigated, transcriptome-wide association analysis was carried out to highlight genes for which a variation in brain gene expression may be causally linked to phenotypes of interest. Our results highlight the role of genomic region 17q21.3 as a locus shared by all four disorders. This region was also identified as an independent genomic risk region from our annotation for SNP-based analyses. The top result of our four-disorder combined transcriptome-wide analysis was from transcript of *LRRC37A4P*. Even though it is a pseudogene, altered transcription levels of this probe have been observed in multiple neurological conditions, including Alzheimer disease and ASD (86–89). Furthermore, the implicated region has been previously found as contributing to various neurodevelopmental or neurodegeneration processes (90–94).

Although we provide results on the largest available combined dataset across ADHD, ASD, OCD, and TS, available datasets varied in size for each of the studied disorders. The unbalanced sample size across the studied datasets is one of the limitations of our study with results sometimes driven by the larger studies and greater difficulty in uncovering loci of importance for under-represented disorders. In order to mitigate this problem, we ran analyses using different pairwise combinations of disorders and we placed emphasis on investigating and reporting the SNP posterior probability of association (m-value) for each disorder in order to allow better interpretation of results and provide higher confidence for shared effect across multiple disorders. Existing overlap across the studied samples was another challenge (< 6% case overlap in the datasets that we studied). In order to tackle this problem, we used ASSET, which takes into account known sample overlap to control the inflation in meta-analysis results.

In conclusion, through a series of systematic genome-wide association meta-analyses we uncovered multiple loci that may underlie biological mechanisms across ADHD, ASD, OCD, and TS. Our results provide further support for the hypothesis of a common etiological basis across this spectrum of often comorbid neurodevelopmental phenotypes. Both the *SORCS3* region and the intergenic region 5q21.2 that were picked up by our study on the four related disorders, were also highlighted as highly pleiotropic loci in the eight-disorders study by the Cross-Disorder Group of the Psychiatric Genomics Consortium (20), indicating a broader functional impact of these regions across multiple psychiatric disorders. However, here we also identify many additional genes and genomic risk loci that could play a more specific role across the spectrum of the ADHD, ASD, OCD, TS phenotypes. The existing evidence for a common genetic background across these highly comorbid disorders highlights what seems to become a recurrent theme across the studies on neuropsychiatric disorders: the importance of thinking across diagnostic boxes when attempting to understand neurobiology. Towards this end, large well-characterized cohorts of patients will be necessary as well as the harmonization of existing clinical databases spanning the disorder spectrum. Analyzing across a spectrum of intermediate phenotypes may hold the promise to identify novel targets for improved therapies focusing on an individual patient rather than a broadly defined diagnostic category.

## Supporting information

Supplementary table 1. SNP-based meta-analyses results

Supplementary table 2. Genomic risk loci

Supplementary table 3. Gene-based analyses results

Supplementary table 4. 53 tissue specificity analyses results

## Supplementary Material

### Supplementary Tables – Legends

**Table S1.** Summary statistics for all significant results from SNP-based GWAS meta-analyses across ADHD, ASD, OCD, TS (four-disorder and pairwise analysis, each in corresponding worksheet). *m-value* = Posterior probability for association for each individual disorder; *SIFT/Poly1/Poly2* = functional prediction for nonsynonymous exonic SNPs; *HetISq* = heterozygosity I^2^ statistic; *HetChiSq* = heterozygosity chi-square statistic; HetPVal = heterozygosity test p-value; *disorder-OR/P* = odds ratio statistic and p-value in the original individual disorder GWAS study.

**Table S2.** Full annotation of top genomic risk regions from SNP-based GWAS meta-analyses. An asterisk (*) indicates novel LD regions not been reported associated with corresponding traits in published GWAS. *rsID* = rsID of the leading SNP of the region; *p* = p-value of the leading SNP from the meta-analysis; *Study* = Previous studies reporting significant association at this locus; *trait* = trait reported associated with the locus by the study; *reported gene* = gene reported by the study; *mapped gene* = gene mapped onto the reported region.

**Table S3.** Significant genes from gene-based GWAS analyses. Also showing p-values from each individual disorder gene-based analyses.

**Table S4.** Significant results from cross-disorder ADHD-ASD-OCD-TS tissue specificity analysis, testing 53 tissue types from GTEx v7 tissue expression atlas. Significant threshold is subjective to Bonferroni correction (p < 9.43 x 10^-4^)

### Supplementary Figures – Legends

**Figure S1.** Top ten gene networks from top 200 genes annotated from SNP-based GWAS meta-analyses results and gene-based analysis results. A. ADHD-ASD SNP-based network plot; B. ADHD-ASD gene-based network plot.

**Figure S2.** Top ten gene networks from top 200 genes annotated from SNP-based GWAS meta-analyses results and gene-based analysis results. A. ADHD-OCD SNP-based network plot; B. ADHD-OCD gene-based network plot.

**Figure S3.** Top ten gene networks from top 200 genes annotated from SNP-based GWAS meta-analyses results and gene-based analysis results. A. ADHD-TS SNP-based network plot; B. ADHD-TS gene-based network plot.

**Figure S4.** Top ten gene networks from top 200 genes annotated from SNP-based GWAS meta-analyses results and gene-based analysis results. A. ASD-TS SNP-based network plot; B. ASD-TS gene-based network plot.

**Figure S5.** Top ten gene networks from top 200 genes annotated from SNP-based GWAS meta-analyses results and gene-based analysis results. A. OCD-TS SNP-based network plot; B. OCD-TS gene-based network plot.

**Figure S6.** ADHD-ASD-OCD-TS cross-disorder tissue specificity analysis, testing 53 tissue types from GTEx v7 tissue expression atlas. Red bar indicates significant enrichment of gene expression in corresponding tissue under Bonferroni correction (p < 9.43 x 10^-4^). Panel on top right corner of each figure shows detailed statistics for significantly enriched tissue. A. ADHD-ASD-OCD-TS cross-disorder tissue specific expression enrichment; B. ADHD-ASD cross-disorder tissue specific expression enrichment; C. ADHD-OCD cross-disorder tissue specific expression enrichment; D. ADHD-TS cross-disorder tissue specific expression enrichment; E. ASD-TS cross-disorder tissue specific expression enrichment; F. OCD-TS cross-disorder tissue specific expression enrichment.

## Acknowledgements

This work was made possible thanks to the concerted efforts of the Psychiatric Genomics Consortium Cross-disorder Working Group, the Lundbeck Foundation Initiative for Integrative Psychiatric Research (iPSYCH), the Psychiatric Genomics Consortium ADHD Working Group, the Psychiatric Genomics Consortium ASD Working Group, and the Psychiatric Genomics Consortium TS/OCD Working Group.

