## Supplementary table legends and supplementary figures for "Cross-disorder GWAS meta-analysis for Attention Deficit/Hyperactivity Disorder, Autism Spectrum Disorder, Obsessive Compulsive Disorder, and Tourette Syndrome"

**Supplementary Material**
**Supplementary Tables – Legends**

**Table S1.** Summary statistics for all significant results from SNP-based GWAS meta-analyses across ADHD, ASD, OCD, TS (four-disorder and pairwise analysis, each in corresponding worksheet). *m-value* = Posterior probability for association for each individual disorder; *SIFT/Poly1/Poly2* = functional prediction for nonsynonymous exonic SNPs; *HetISq* = heterozygosity $I^2$ statistic; *HetChiSq* = heterozygosity chi-square statistic; HetPVal = heterozygosity test p-value; *disorder-OR/P* = odds ratio statistic and p-value in the original individual disorder GWAS study.

**Table S2.** Full annotation of top genomic risk regions from SNP-based GWAS meta-analyses. An asterisk (*) indicates novel LD regions not been reported associated with corresponding traits in published GWAS. *rsID* = rsID of the leading SNP of the region; *p* = p-value of the leading SNP from the meta-analysis; *Study* = Previous studies reporting significant association at this locus; *trait* = trait reported associated with the locus by the study; *reported gene* = gene reported by the study; *mapped gene* = gene mapped onto the reported region.

**Table S3.** Significant genes from gene-based GWAS analyses. Also showing p-values from each individual disorder gene-based analyses.

**Table S4.** Significant results from cross-disorder ADHD-ASD-OCD-TS tissue specificity analysis, testing 53 tissue types from GTEx v7 tissue expression atlas. Significant threshold is subjective to Bonferroni correction ($p < 9.43 \times 10^{-4}$)

**Figure S1**. Top ten gene networks from top 200 genes annotated from SNP-based GWAS meta-analyses results and gene-based analysis results.  A. ADHD-ASD SNP-based network plot; B. ADHD-ASD gene-based network plot.

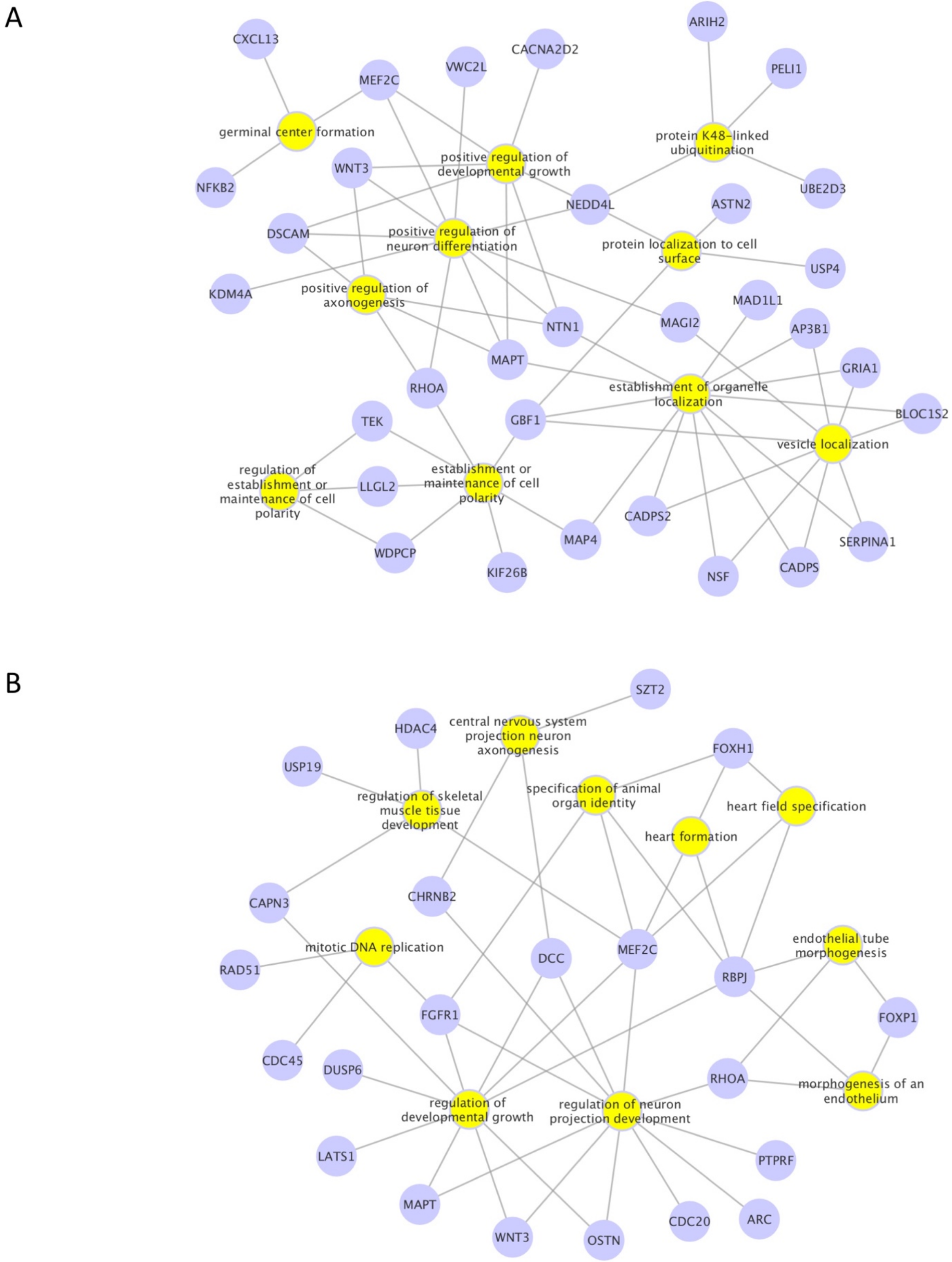

**Figure S2**. Top ten gene networks from top 200 genes annotated from SNP-based GWAS meta-analyses results and gene-based analysis results. A. ADHD-OCD SNP-based network plot; B. ADHD-OCD gene-based network plot.

A

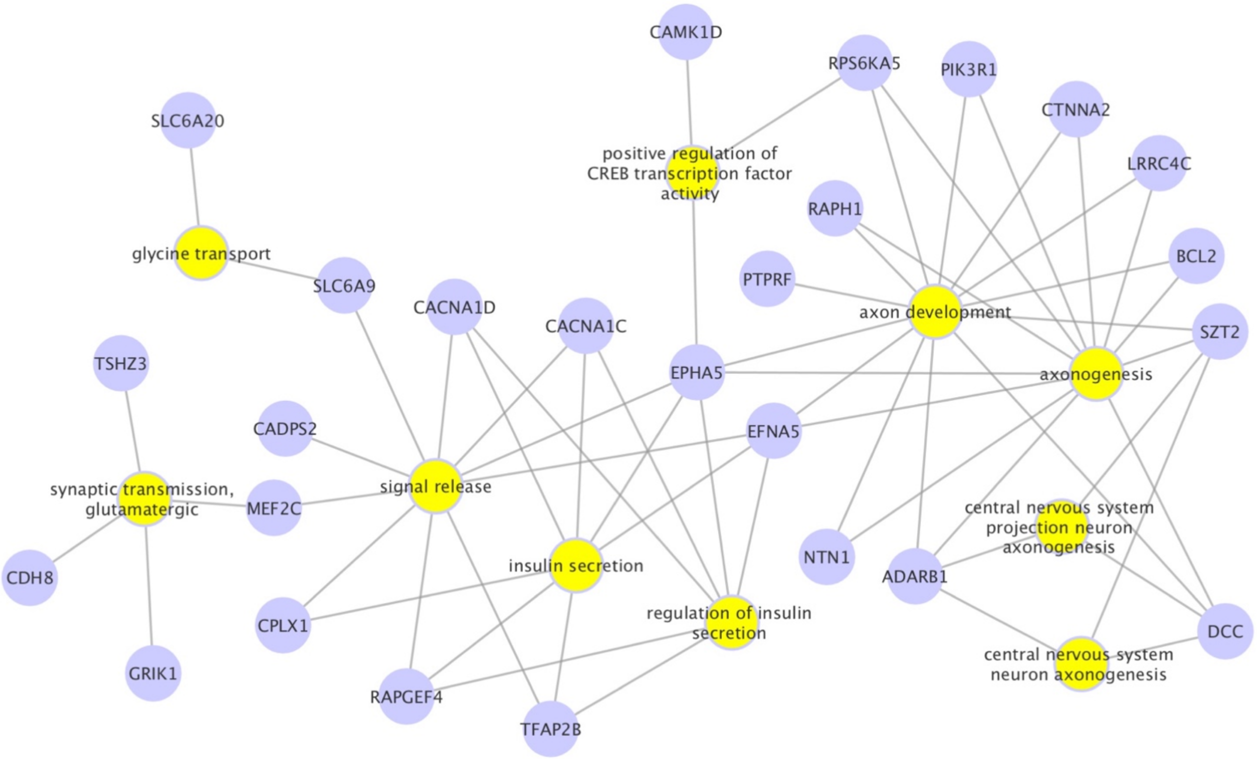


B

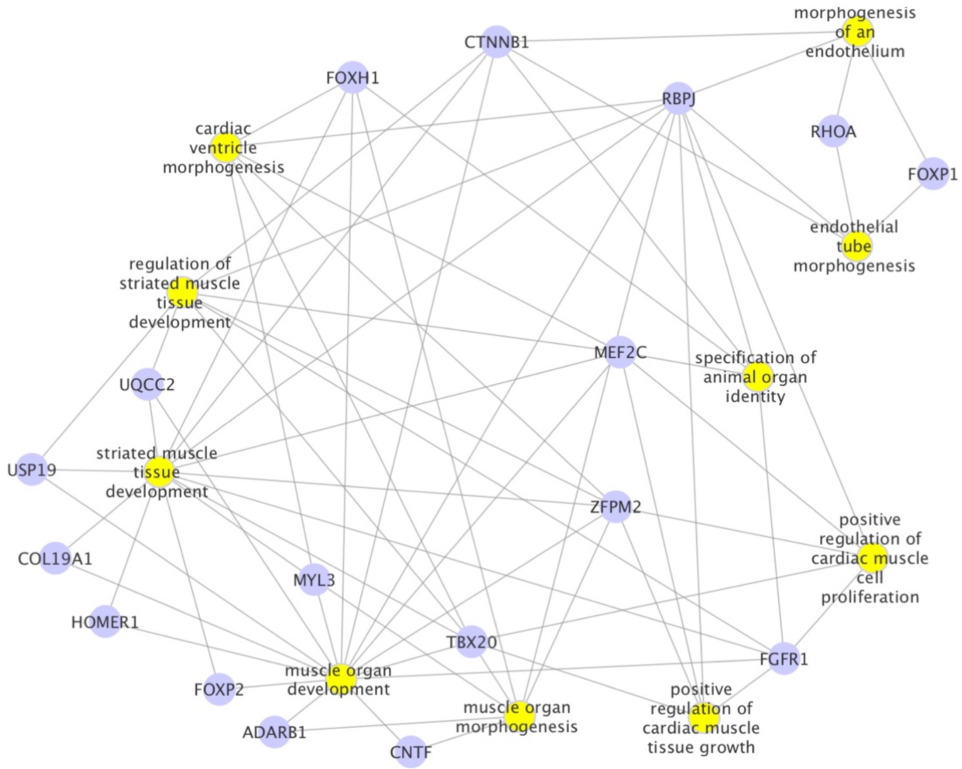

**Figure S3**. Top ten gene networks from top 200 genes annotated from SNP-based GWAS meta-analyses results and gene-based analysis results. A. ADHD-TS SNP-based network plot; B. ADHD-TS gene-based network plot.

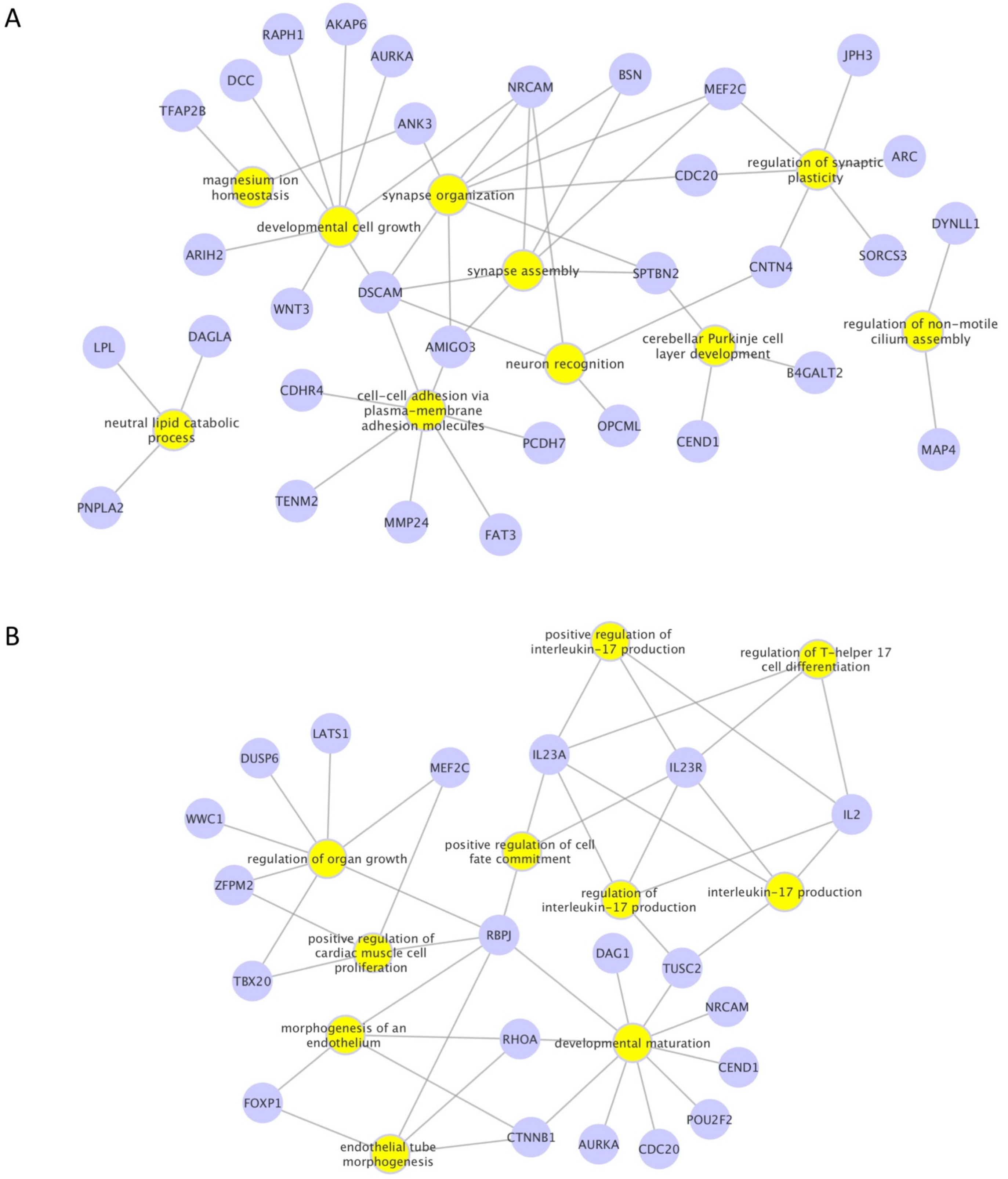

**Figure S4**. Top ten gene networks from top 200 genes annotated from SNP-based GWAS meta-analyses results and gene-based analysis results. A. ASD-TS SNP-based network plot; B. ASD-TS gene-based network plot.

A

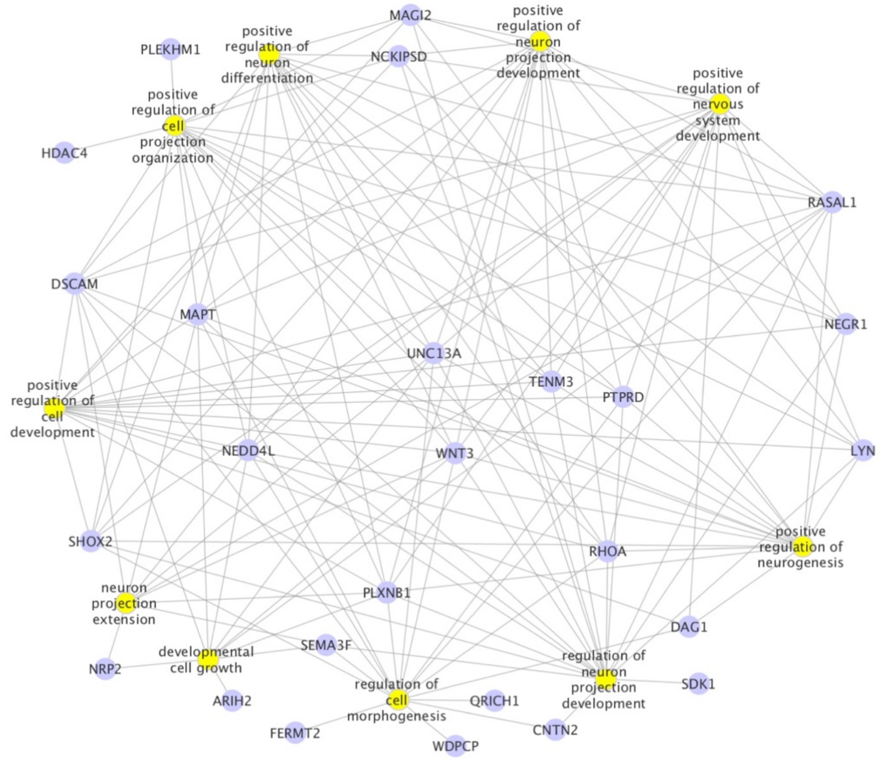


B

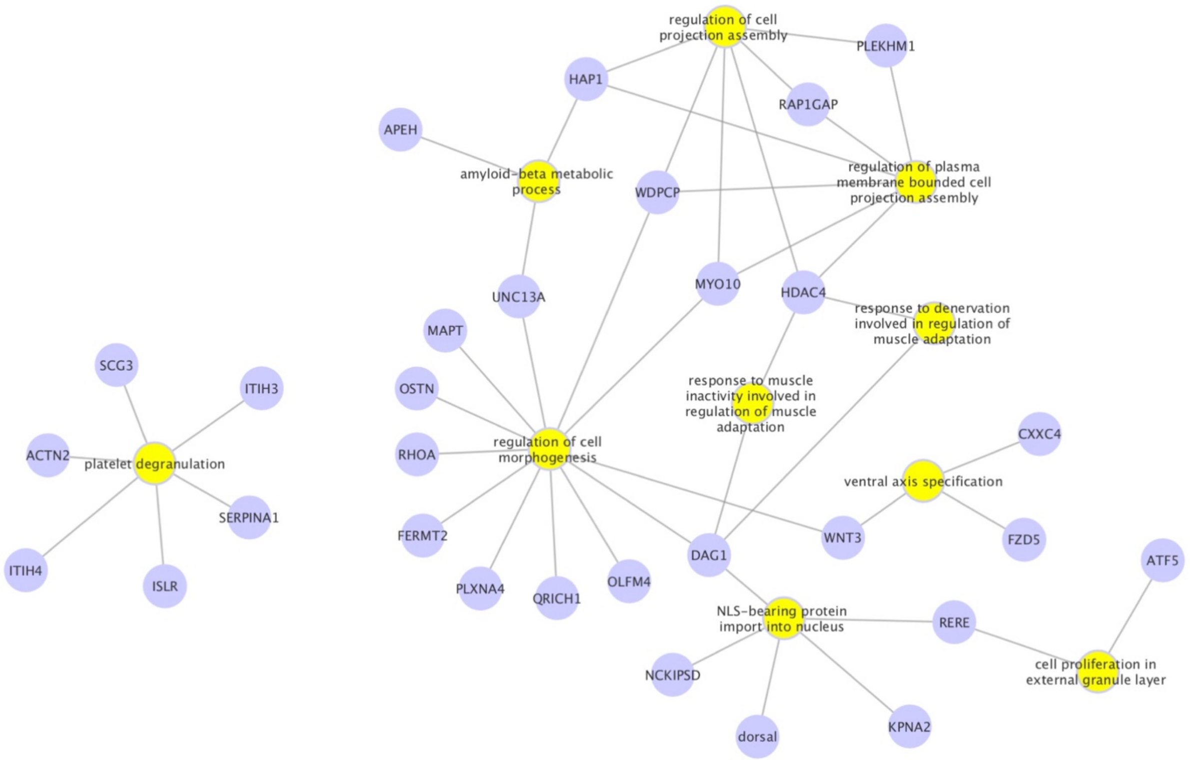

**Figure S5**. Top ten gene networks from top 200 genes annotated from SNP-based GWAS meta-analyses results and gene-based analysis results. A. OCD-TS SNP-based network plot; B. OCD-TS gene-based network plot.

A

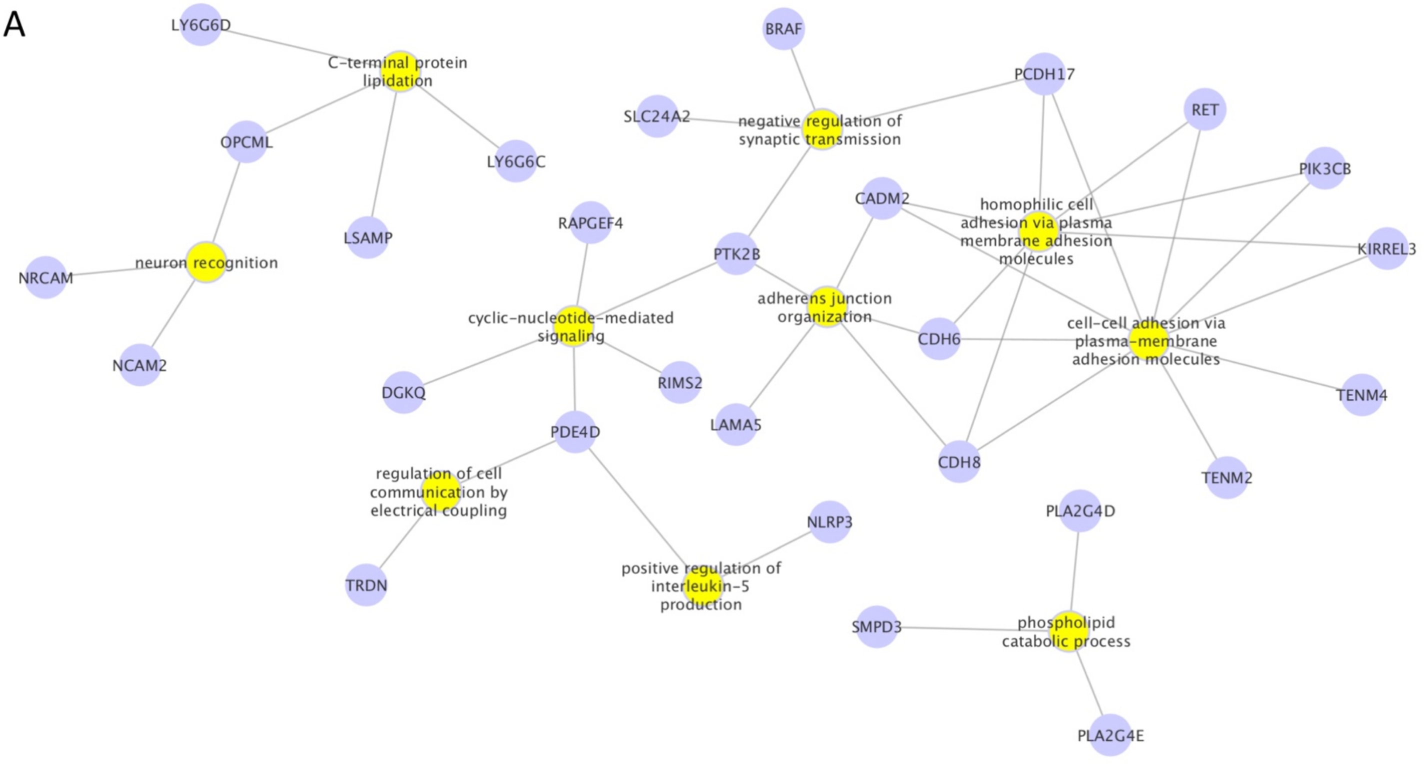


B

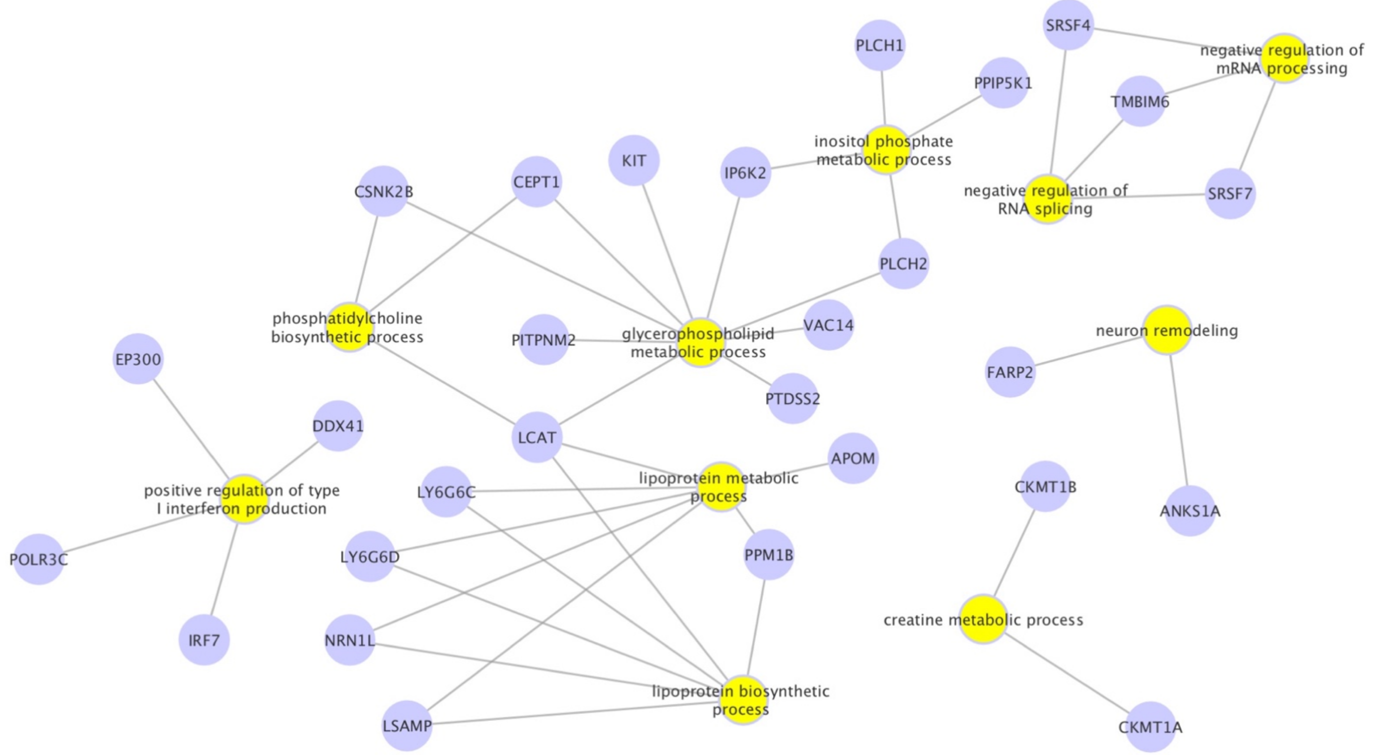

**Figure S6**. ADHD-ASD-OCD-TS cross-disorder tissue specificity analysis, testing 53 tissue types from GTEx v7 tissue expression atlas. Red bar indicates significant enrichment of gene expression in corresponding tissue under Bonferroni correction ($p < 9.43 \times 10^{-4}$). Panel on top right corner of each figure shows detailed statistics for significantly enriched tissue. A. ADHD-ASD-OCD-TS cross-disorder tissue specific expression enrichment; B. ADHD-ASD cross-disorder tissue specific expression enrichment; C. ADHD-OCD cross-disorder tissue specific expression enrichment; D. ADHD-TS cross-disorder tissue specific expression enrichment; E. ASD-TS cross-disorder tissue specific expression enrichment; F. OCD-TS cross-disorder tissue specific expression enrichment

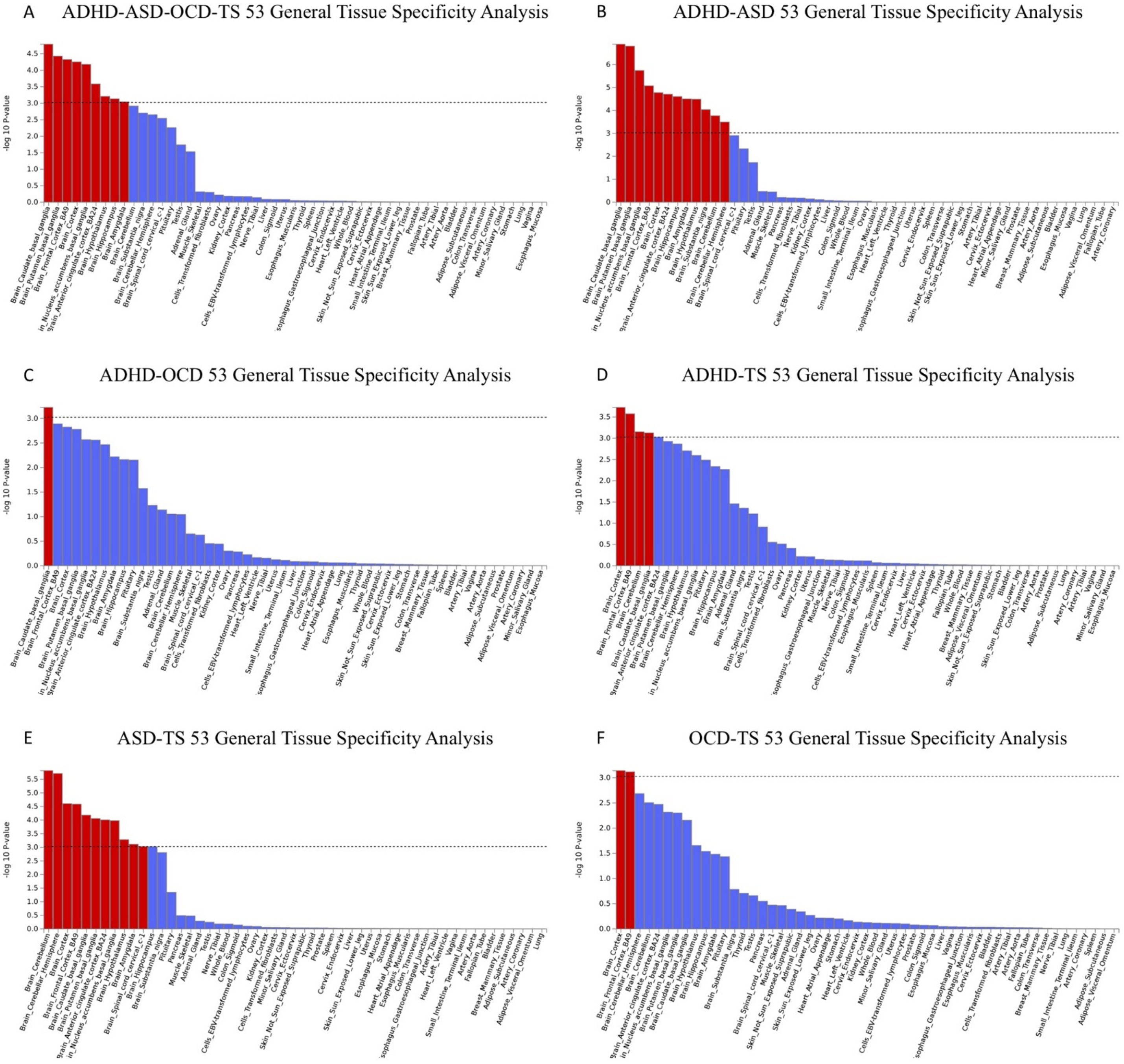


.
